# Analysis of CHD-7 defective dauer nematodes implicates collagen misregulation in CHARGE syndrome features

**DOI:** 10.1101/2021.03.26.437191

**Authors:** Diego M. Jofré, Dane K. Hoffman, Ailen S. Cervino, McKenzie Grundy, Sijung Yun, Francis RG. Amrit, Donna B. Stolz, Esteban Salvatore, Fabiana A. Rossi, Arjumand Ghazi, M. Cecilia Cirio, Judith L. Yanowitz, Daniel Hochbaum

**Author notes:** Authors contributed equally (Diego M. Jofré, Dane K. Hoffman). Department of Cancer & Cell Biology. Baylor College of Medicine. Houston, Texas, USA. Co-corresponding authors. **Correspondence:** Correspondence should be addressed to Dr. Daniel Hochbaum, Ciudad Universitaria. Intendente Guiraldes 2160, 4 piso. Laboratorio 79, CABA, Buenos Aires., C1428BGA, Argentina, Dr. Judith L. Yanowitz, Magee-Womens Research Institute, 204 Craft Avenue, Pittsburgh, PA, 15213, United States of America. **Author Contributions** Conceived and designed the experiments: DMJ, DKH, ASC, FAG, ES, MCC, JLY, DH. Performed the experiments: DMJ, DKH, ASC, MG, FRGA, DBS, ES, FAR, DH. Analyzed the data: DMJ, DKH, ASC, SY, FRGA, AG, MCC, JLY, DH. Drafted the manuscript: MCC, JLY, DH. Reviewed and edited the manuscript: DMJ, DKH, AG, MCC, JLY, DH.

## Abstract

CHARGE syndrome is a complex developmental disorder caused by mutations in the chromodomain helicase DNA-binding protein7 (CHD7) and characterized by retarded growth and malformations in the heart and nervous system. Despite the public health relevance of this disorder, relevant targets of CHD7 that relate to disease pathology are still poorly understood. Here we report that *chd-7*, the nematode ortholog of Chd7, is required for dauer morphogenesis, lifespan determination, and stress response. Consistent with our discoveries, we found *chd-7* to be allelic to *scd-3*, a previously identified dauer suppressor from the TGF-β pathway. Notably, DAF-12 promoted *chd-*7 expression, which is necessary to repress *daf-9* for execution of the dauer program. Transcriptomic analysis comparing *chd-7–*defective and normal dauers showed enrichment of collagen genes, consistent with a conserved role for the TGF-β pathway in formation of the extracellular matrix. To validate a conserved function for *chd-7* in vertebrates, we used *Xenopus laevis* embryos, an established model to study craniofacial development. Morpholino mediated knockdown of Chd7 led to a reduction in *col2a1* mRNA levels. Both embryonic lethality and craniofacial defects in Chd7-depleted tadpoles were partially rescued by over-expression of *col2a1* mRNA. We suggest that pathogenic features of CHARGE syndrome caused by Chd7 mutations, such as craniofacial malformations, result from the reduction of collagen levels, implying that the extracellular matrix might represent a critical target of Chd7 in CHARGE development.

**SIGNIFICANCE STATEMENT:** CHARGE Syndrome is a complex developmental disorder caused by mutations in the chromodomain helicase DNA-binding protein-7 (CHD7). Unfortunately, the cellular events that lead to CHARGE syndrome are still poorly understood. In *C. elegans*, we identified *chd-7* in a screen for suppressors of dauer formation, an alternative larval stage that develops in response to sensory signals of a harsh environment. We found that *chd-7* regulates expression of collagens, which constitute the worm’s cuticle, a specialized extracellular matrix. In frog’s embryos, we show that Chd7 inhibition leads to poor Col2a1, which is necessary and sufficient to exhibit CHARGE features. These studies establish *C. elegans* as an amenable animal model to study the etiology of the developmental defects associated with pathogenic Chd7.

## INTRODUCTION

CHARGE syndrome is a rare and severe neurodevelopmental disorder that affects the neural tube and neural crest cell derivatives, leading to hypogonadism, heart defects and craniofacial anomalies among other features (1). Inactivating mutations in *CHD7* (chromodomain-helicase-DNA binding 7) are the predominant cause of CHARGE, accounting for greater than 90% of the cases (2). *CHD7* is also mutated in Kallmann syndrome, a milder neurodevelopmental disorder with features overlapping with CHARGE, including impaired olfaction and hypogonadism (3). Exome sequencing studies in patients with autism spectrum disorders (ASDs) identified recurrent disruptive mutations in the related gene *CHD8* (4). The CHD proteins comprise a highly conserved family of SNF2-related ATP-dependent chromatin remodelers that are involved in chromatin remodeling and transcriptional regulation (5). Despite the public health relevance of these cognitive disorders, the mechanism of disease pathology due to mutations in CHD7/8 is poorly understood. The development of fly, fish and mouse models of CHARGE has abetted in characterization of associated dysfunction, but our understanding of the underlying pathology of CHARGE is still incomplete (6–11). CHD-7 is the ortholog of human CHD7 and CHD8 in *Caenorhabditis elegans* and has functions in habituation learning, normal locomotion, body size and fecundity (12, 13). It contains a conserved ATPase/SNF2 domain and two chromodomains for nucleosome interaction. Being the only worm homolog of the Class III CHD family, it contains a signature BRK domain (Brahma and Kismet domain) (Figure 1E).

**Figure 1:**
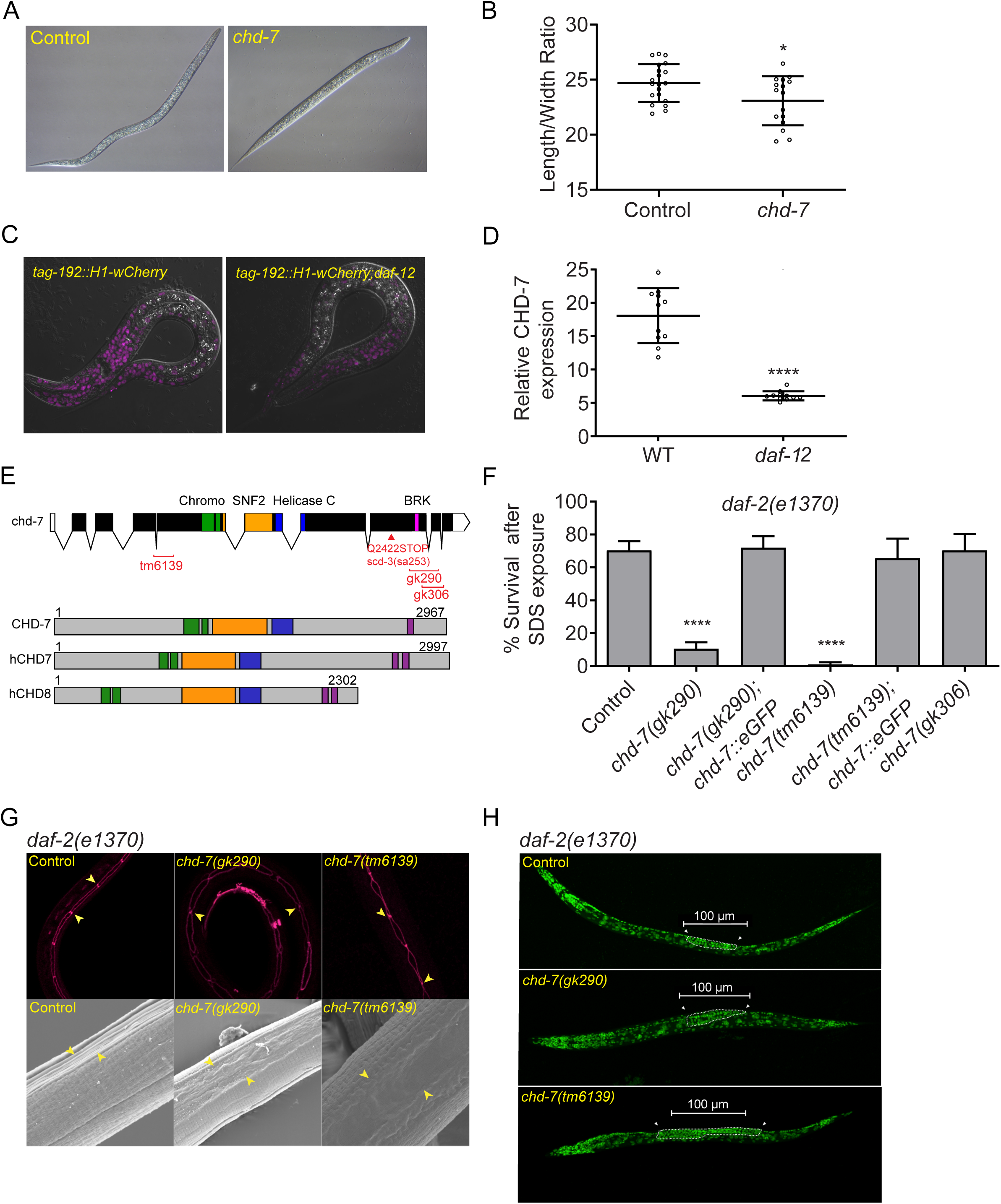
The DAF-12 regulated target *chd-7* is required for proper dauer morphogenesis. A) *chd-7(RNAi)* causes a partial dauer phenotype in *daf-2(e1370).* Representative DIC photomicrographs of normal and partial dauers from *daf-2(e1370)* exposed to Control (L4440) or *chd-7* dsRNA, respectively. B) Quantification of axial ratio of *daf-2(e1370);*Control(RNAi) and *daf-2(e1370);chd-7(RNAi)* dauers. Three biological replicates were scored (n=16-21/replicate). Horizontal black lines represent mean with SD. Unpaired t test, *p<0.05. C) *daf-12* regulates *chd-7* expression. Representative images of *chd-7* transcriptional reporter, *tag-192::H1-wCherry* or *tag-192::H1-wCherry;daf-12(rh61rh411)* worms at L2/L3 stage. D) Relative expression of the translational reporter (n>10/strain). Unpaired t test, ****p<0.0001. E) *C. elegans chd-*7 gene and protein. Top: *chd-7* genomic region. UTR and exons shown as bars; introns by lines. In red, available *chd-7* deletional alleles (data obtained from CGC and NBRP). Bottom images depict the predicted protein isoforms of *C. elegans* CHD-7, human CHD7 and human CHD8. Signature domains in CHD proteins: two N-terminal chromodomains for interaction with a variety of chromatin components (green), a SNF-2 like domain with ATPase activity (yellow) and a Helicase domain (blue). The Class III subfamily is defined by a BRK domain (purple). F) *chd-7*(*gk290*);*daf-2(e1370)* and *chd-7*(*tm6139*);*daf-2(e1370)* develop as SDS-sensitive dauer larvae. n>725 animals/strain tested. Bars and horizontal black lines represent mean percentage with SD. Chi-squared test with Bonferroni correction for multiple comparisons. ****p<0.0001. * represents the comparison to the *daf-2(e1370)* strain G) *chd-7* partial dauers fail to develop the dauer alae. Top row, representative photomicrographs of *daf-2(e1370)* dauers or *chd-7;daf-2(e1370)* partial dauers expressing the *ajm-1*::GFP reporter to delineate the seam cell borders (arrowheads mark a subset of junctions). Bottom row, SEM images of *daf-2(e1370)* dauers or *chd-7;daf-2(e1370)* partial dauers (arrowheads mark alae details). H) Germ cells in *chd-7;daf-2(e1370)* mutants overproliferate and arrest as L3-like germ cells. Representative Z-projections *daf-2(e1370)* dauers or *chd-7;daf-2(e1370)* partial dauers stained with DAPI (green). Arrowheads denote the anterior and posterior ends of the developing gonad arms.

When *C. elegans* encounter crowding, starvation or high temperature during early development, worms can halt reproductive programs to enter an alternative larval stage known as dauer. Dauers are long-lived, highly stress resistant and exhibit altered motility and metabolism (14–17). Upon return to normal growth conditions, the larvae exit dauer and develop into fertile adults. Study of dauer formation mutants has provided fundamental insights into pathways affecting longevity, neurodevelopment, metabolism, autophagy and neurodegeneration (14, 17–20).

The DAF-2/insulin/IGF-signaling 1 (IIS) pathway controls the dauer entry decision by coupling external cues with neuroendocrine signaling (21). In favorable conditions, DAF-2 activity initiates a conserved kinase cascade, leading to phosphorylation and inhibition of the transcription factor DAF-16/FOXO. In harsh environments, a decrease in the activity of DAF-2 and downstream components of the pathway leads to activation of DAF-16 and causes animals to arrest as dauers (22, 23). In addition to the DAF-2 pathway, DAF-7/ TGF-β signaling also regulates dauer development (24). When worms sense suitable conditions for reproductive development, ASI neurosensory cells secrete DAF-7, which binds to DAF-1/4 receptors leading to activation and phosphorylation of the R-SMAD complex DAF-8/14, promoting reproductive programs and inhibiting the pro-dauer complex composed of the SMAD protein DAF-3 and repressor DAF-5. Conversely, absence of DAF-7 leads to activation of the DAF-3/DAF-5 complex to promote dauer entry (25, 26).

DAF-7/TGF-β and DAF-2/IIS pathway were initially described as parallel pathways to regulate dauer entry (27), but recent observations suggest a strong positive feedback between these pathways for dauer entry and longevity (15–20). First, decreased signaling through the TGF-β pathway leads to differential expression of many DAF-16 regulated genes with functions in longevity and dauer entry, such as SOD-3 or insulin peptides. This cross-activation of target genes may be important to amplify weak signals from each sensory pathway to make an all-or-none decision to enter dauer (28–30). Second, the longevity of *daf-2* mutants can be blocked or enhanced by *daf-5* and *daf-3*, respectively, suggesting that transcriptional components of the TGF-β can modulate IIS-dependent longevity genes. Third, for dauer development, *daf-16* is epistatic to *daf-7/8/14 daf-c* mutants (30, 31). Lastly, both signaling pathways converge on *daf-9* and *daf-12* to integrate outputs for diapause entry (32, 33). Indeed, *daf-9* expression levels is critical for both entering and exiting diapause (17, 32).

In chromatin immunoprecipitation (ChIP)-chip studies, we identified *chd-*7 as a DAF-12 target gene whose loss caused defective execution of dauer morphogenesis programs (34). We show that while CHD-7 can modulate multiple IIS-associated processes, including *daf-2(e1370)* dauer formation, longevity and immunity, epistatic experiments place *chd-7* in the DAF-7/TGF-β pathway. Whole genome mRNA expression profiling of partial dauers show that *chd-7* mutants fail to repress *daf-9* during dauer development, preventing developmental arrest of the constitutive dauer mutant *daf-7(e1372)*. In addition, we found collagens to be among the most differentially regulated genes, consistent with a known role for the TGF-β pathway in regulating components of the extracellular matrix (35).To validate a conserved function for *chd-7* in vertebrates and study the relevance of our results for CHARGE etiology, we used *Xenopus laevis*. Disruption of Chd7 function in *Xenopus* embryos results in craniofacial defects that mimic CHD-dependent pathological phenotypes (36, 37). We demonstrate that Chd7 regulates expression of the collagen type-II, alpha 1, *col2a1*, the main collagen protein of cartilage (38). Interestingly, craniofacial malformations and embryonic lethality due to *chd7* knockdown can be rescued by *col2a1* expression. These findings suggest a conserved function of *chd7*/*chd-7* in regulation of extracellular matrix components and raise the intriguing possibility that defects in collagen expression may contribute to the craniofacial defects seen in CHD7/8-dependent syndromes.

## RESULTS

### *chd-7* functions in development of the dauer larva

Previously, we identified ∼3000 potential DAF-12 target genes by ChIP-chip (34). We reasoned that among these genes would be novel regulators of dauer development. To identify such genes, we took advantage of the temperature-sensitive, constitutive dauer (Daf-c) allele *daf-2(e1371)*. At the non-permissive temperature of 25°C, all the offspring enter the dauer phase and become SDS resistant (22). We expected that inactivating targets of DAF-12 that function in dauer formation should suppress the Daf-c phenotype of *daf-*2 and produce either defective, SDS-sensitive “partial” dauers or reproductive adults. To confirm these results, we assayed suppression of the more severe *daf-2(e1370)* mutation and discovered that *chd-7(RNAi)* produced arrested, SDS-sensitive larvae, indicative of the partial dauer phenotype (39). As observed in Figure 1A and B, the axial ratio (length/width) of the partial dauers resulting from *chd-7(RNAi)* exhibited a significant reduction of these proportions and appeared to have defects in radial constriction of the dauer cuticle. By using a *chd-7* transcriptional reporter (WBStrain00033709), we observed a substantial decrease in *chd-7* expression in the *daf-12(rh61rh411)* background (Figure 1C and D). Thus, we infer that DAF-12 binds to the *chd-7* promoter (34) and upregulates its expression.

To further validate our screen, we crossed *daf-2(e1370)* mutants with three *chd-7* deletion alleles available from the Nematode Knockout Consortia (40). Allele *chd-7(tm6139)* contains a 594bp deletion that generates a frame shift and premature stop codon, eliminating all known protein domains (Figure 1E). As shown in Figure 1F, SDS-sensitive dauers were obtained when double mutants are grown at 25°C, validating our screen. Comparison of two partial deletion alleles uncovered a critical role for the BRK domain in dauer formation. The *chd-7(gk290)* allele contains a 859bp deletion that spans the BRK domain and introduces a frameshift that eliminates the last 356aa. The *chd-7(gk306)* deletion is slightly more C-terminal, truncating the protein immediately after the BRK domain (Figure 1E). When crossed into *daf-2(e1370)* worms, *chd-7(gk306)* developed normal, SDS-resistant dauer larvae, whereas *chd-7(gk290)* formed partial dauers that were sensitive to detergent (Figure 1F). Importantly, a functional transgene expressing GFP-tagged CHD-7 protein (CHD-7::GFP) rescued the partial dauer phenotypes observed in *chd-7(gk290);daf-2(e1370)* and *chd-7(tm6139);daf-2(e1370)* mutants (Figure 1F).

The dauer larva is characterized by a slim physique because of a reduction in the volume of ectodermal tissues, including the hypodermis, seam cells and pharyngeal cells. In addition, the hypodermis produces the dauer cuticle, which confers protection against external damage and dehydration. During development, the seam cells fuse and the multinucleate cells produce the alae-bilateral ridges in the cuticle that facilitate body motion. Dauer larvae also switch their metabolism to accumulate lipids to survive for longer periods. To further characterize the nature of the defects in the *chd-7-*induced partial dauers, we first studied the seam cells fusions with the adherens junction-associated protein marker AJM-1::GFP and the morphology of the cuticle with scanning electron microscopy (SEM). In dauering preparation, animals store fat in their intestinal and hypodermal cells, which is critical to survive during hibernation (41). As shown in Figure 1G, partial dauers exhibited defects in seam cell fusion and defective dauer alae formation. Next we used the lipid-labeling dye, Oil Red O (ORO), to examine fat storage and found that, unlike other partial dauer mutants (19), the *chd-7;daf-2* abnormal dauers did not exhibit fat-storage deficiencies (Supplemental Figure 1).

During dauer, the global developmental arrest also impacts the germ cells, slowing their divisions and finally remaining quiescent (42). We observed that the germ line in *chd-7* mutant dauers was substantially larger than in control dauers and appeared to arrest with a germline morphology resembling that seen in L3 larval stage (Figure 1H). Therefore, we conclude that major morphological changes that occur during dauer formation fail to be executed in *chd-7* mutants, including radial constriction of the body, formation of an SDS-resistant cuticle with dauer alae, and developmental arrest of the germ line.

### *chd-7* is required for longevity and immunoresistance induced by IIS inactivation and germline removal

In addition to dauer development, the IIS pathway also regulates longevity (14). Hence, we sought to investigate whether *chd-7* also has roles in the determination of lifespan. First, we compared survival of wild-type (N2) worms with two *chd-7* alleles and found that the dauer-defective allele *chd-7(gk290)* significantly shortened lifespan, whereas *chd-7(gk306)* had only a marginal effect on longevity (Figure 2A). Surprisingly, the CHD-7::GFP rescue transgene also reduced N2 lifespan (Figure 2B), suggesting that *chd-7* copy number can influence longevity. We then analyzed how *chd-7* affects longevity of IIS mutants. Remarkably, the dauer suppressor allele *chd-7(gk290),* but not *chd-7(gk306),* shortened the lifespan extension of *daf-2* mutants to an extent comparable with the null allele of *daf-16,* the key IIS downstream target (Figure 2C) (14). Furthermore, *daf-2* longevity was fully restored by CHD-7::GFP (Figure 2D).

**Figure 2:**
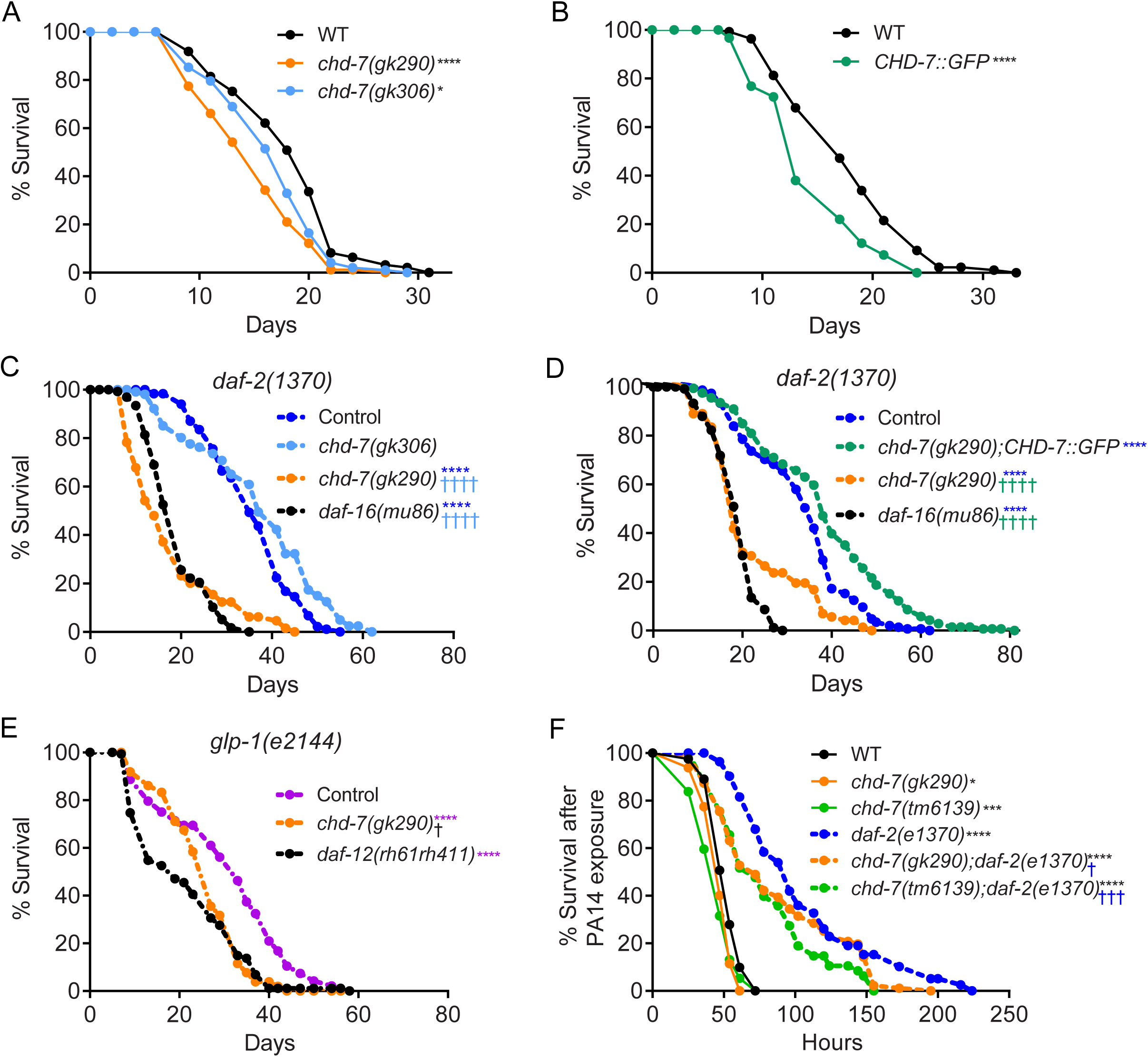
*chd-7* affects longevity and response to pathogen. A-E) *chd-7* promotes longevity in wild-type, *daf-2(e1370)* and *glp-1(e2144)* mutants. Mean survival days on OP50-1; survival data analyzed using Kaplan−Meier test. A) WT (18.12 days), *chd-7(gk290*) (14.91), *chd-7(gk306)* (16.53). *p<0.05 and ****p<0.0001 compared to the wild-type, N2 strain. B) WT (17.85), CHD-7::GFP (14.35). ****p<0.0001 compared to the wild-type, N2 strain. C) *daf-2(e1370)* (35.58)*, chd-7(gk306);daf-2(e1370)* (37.18)*, chd-7(gk290);daf-2(e1370)* (17.47)*, daf-16(mu86);daf-2(e1370)* (18.85). ****,^††††^p<0.0001. *represents comparison to *daf-2(e1370)*; ^†^represents comparison to *chd-7(gk306);daf-2(e1370)*. D) *daf-2(e1370)* (27.9)*, chd-7(gk290);daf-2(e1370);CHD-7::GFP* (30.84)*, chd-7(gk290);daf-2(e1370)* (19.13)*, daf-16(mu86);daf-2(e1370)* (14.48). ****,^††††^p<0.0001. *represents comparison to *daf-2(e1370);* ^†^represents comparison to *chd-7(gk290);daf-2(e1370);CHD-7::GFP*. E) *glp-1(e2144)* (30.4), *chd-7(gk290);glp-1(e2144)* (28.37), *glp-1(e2144);daf-12(rh61rh411)* (25.99). ****p<0.0001 and ^†^p<0.05. *comparison to *glp-1(e2144)*; ^†^comparison to *glp-1(e2144);daf-12(rh61rh411)*. Details of number of animals and other data from replicates found in Supplemental Table 2. F) *chd-7* mediates the response against the opportunistic bacteria *P. aeruginos*a. Mean lifespan in hours (m) ± standard error of the mean (SEM). ‘n’ is the number of animals analyzed/total number in experiment. WT (m = 52.11 ± 0.96, n = 94/162), *chd-7(gk290)* (m = 47.45 ± 1.05, n = 54.130/), *chd-7(tm6139)* (m = 44.47 ± 1.57, n = 56/80), *daf-2(e1370)* (m = 105.46 ± 5.61, n = 65/115), *chd-7(gk290);daf-2(e1370)* (m = 86.74 ± 4.21, n = 103/120) and *chd-7(tm6139);daf-2(e1370)* (m = 79.13 ± 4.9, n = 50/60). *,^†^p<0.05; ***,^†††^p <0.001 and ****p <0.0001. *comparison to N2; ^†^comparison to *daf-2(e1370)*. Survival data analyzed using Kaplan−Meier test.

To determine if the effects on lifespan were specific to the IIS pathway, we assayed the contribution of *chd-7* to the longevity induced by germ cell-less mutations, a longevity paradigm that operates in parallel to IIS. Temperature-sensitive *glp-1(e2144)* animals are sterile and long-lived at non-permissive temperatures (43). This lifespan extension was dependent on *chd-7*, as *chd-7(gk290);glp-1(e2144)* double mutants had a mean lifespan significantly shorter than *glp-1(e2144)* single mutants (Figure 2E). The impact was similar to that produced by absence of the nuclear receptor *daf-12* (Figure 2C), which is strictly necessary for germless longevity (43).

In addition to longevity, IIS reduction also enhances resistance against multiple stressors including pathogen attack and starvation. We asked if *chd-7* inactivation impaired *daf-2* immunoresistance too. As shown in Figure 2F, *chd-7* mutants repressed the increased survival of *daf-2* worms upon exposure to the human opportunistic pathogen, *Pseudomonas aeruginosa* strain PA14 (44). In contrast, *chd-7* had a modest effect on survival of wild-type worms’ pathogen resistance. Furthermore, the *chd-7(gk290)* and *chd-7(tm6139)* alleles reduced the survival of *daf-2(e1370)* L1 larvae subjected to starvation stress (Supplemental Figure 2) (45). Taken together, these results suggest that *chd-7* mediates the increased lifespan of at least two longevity paradigms (IIS mutants and germ cell-less animals), as well as the response to pathogens and starvation.

### CHD-7 is a member of the TGF-β pathway

Since our genetic analyses implicated *chd-7* as a dauer defective mutant (Daf-d) in the IIS signaling pathway (Figure 1), we sought to understand whether it also had roles in the TGF-β dauer pathway (27, 28). As shown in Figure 3A, at the restrictive temperature of 25°C, *chd-7*;*daf-7(e1372)* double mutants bypassed the dauer arrest to become fertile adults. For *daf-5*, which acts as a dauer suppressor of the TGF-β pathway, the Daf-d phenotype is observed at 25°C, but at a slightly higher temperature, suppression is incomplete (26). Thus, we asked whether *chd-7* also prevented dauer arrest at higher temperatures. As shown in Figure 3A, *daf-7*-dependent arrest was recapitulated at 26.5°C, suggesting that *chd-7* has a Hid (high temperature-induced dauer formation) phenotype (46). As expected for a putative transcriptional regulator, epistasis analysis placed CHD-7 downstream of the receptor DAF-1 and the R-Smad DAF-14 (Figure 3B).

**Figure 3:**
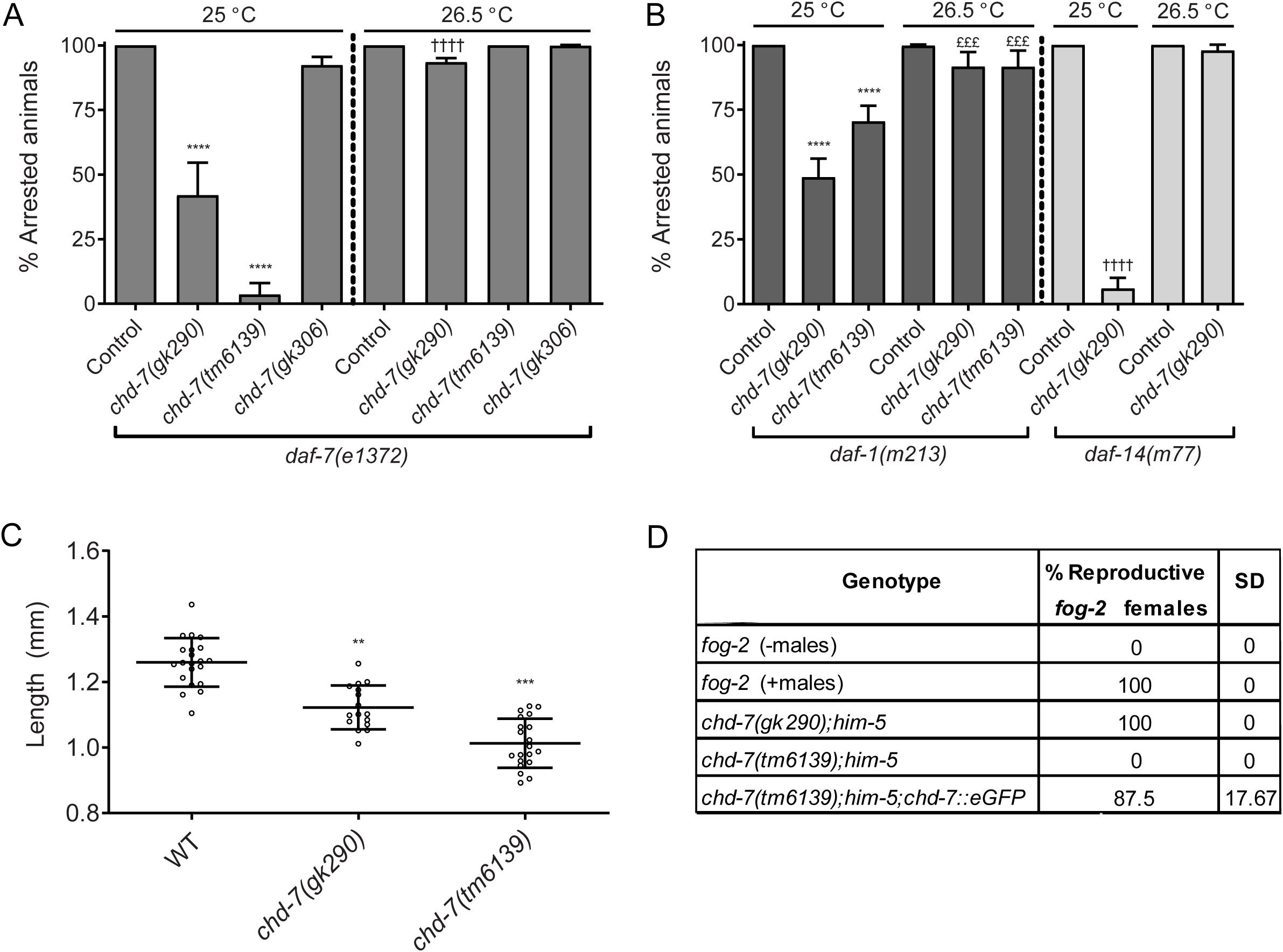
CHD-7 belongs to the TGF-β signaling pathway. A-B) Loss of *chd-7* suppresses dauer arrest of TGF-β pathway mutants at 25°C but not 26.5°C. 7 L4’s were plated individually and grown at the specified temperature for 1 week when arrested and non-arrested progeny were scored. Bars and horizontal black lines represent mean percentage with SD. Statistical significance was calculated using Chi-squared test with Bonferroni correction for multiple comparisons. ****,^††††^p<0.0001 and £££ p <0.001. A) Quantification of dauer arrest in *chd-7;daf-7(e1372)* mutants. * comparison to *daf-7(e1372)* grown at 25°C and ^†^comparison to *daf-7(e1372)* at 26.5°C. B) Dauer arrest in TGF-β pathway mutant backgrounds. *comparison to *daf-1(m213)* grown at 25°C, ^†^comparison to *daf-14(m77)* at 25°C; ^£^comparison to *daf-1(m213)* at 26.5°C. C) *chd-7* regulates body size. Body length of day 1 adults at 20°C (n>16). One-way Anova, **p<0.01, ***p<0.001 compared to wild-type, N2 strain. D) *chd-7(tm6139)* males do not mate. 8 males from each strain tested were plated with 4 *fog-2* females on 10 cm plates. After 24 hrs, *fog-2* females were transferred to new plates and within 48h the proportion of fertile females were scored. Assay was repeated twice.

In addition to dauer formation, the TGF-β signaling pathway also regulates body size and male tale development, mainly through the ligand DBL-1 (47, 48). To test if *chd-7* impacted these processes as well, we measured the length of *chd-7* young adults and found them to be significantly shorter than N2 control animals (Figure 3C). We also observed that males carrying the severe loss-of-function allele *chd-7(tm6139)* failed to mate with *fog-2* mutant females (Figure 3D) due to defects in male tail development (not shown). This male infertility was rescued by the CHD-7::GFP transgene (Figure 3D).

More than twenty years ago, in a screen for suppressors of dauer formation within the TGF-β pathway, *Inoue et al.* identified three complementation groups (31). We noticed that one of these, *scd-3* (<u>s</u>uppressor of <u>c</u>onstitutive <u>d</u>auer-3), was located between *unc-11* and *dpy-5* on chromosome I, in the same genetic region as *chd-7*. Features of *scd-3(sa253)* worms include low brood size, egg-laying defects (Egl), short body size (Dpy) and male abnormal defects (Mab), all of which were phenotypes shared with *chd-7* as described above. In addition, improper gonad migration is a common phenotype of *scd-3* and *chd-7* mutants (Supplemental Figure 4). To determine if *chd-7* and *scd-3* are allelic, we sequenced *scd-3(sa253)* and found a single G/A mutation in exon 8 of the *chd-7* locus, that introduces a premature STOP codon at position Q2422, eliminating the BRK domain of CHD-7 protein, confirming that *chd-7* is *scd-3* (Figure 1E).

### Transcriptomics analysis identifies *daf-9* and collagens as targets of CHD-7

To gain further insight into the role of CHD-7 in dauer development, we performed RNA-Seq. We compared the transcriptomes of *daf-2(e1370)* dauers and *chd-7(gk290);daf-2(e1370)* partial dauers. Differentially expressed genes (DEGs) analysis revealed that decreased expression of 28 genes and increased expression of 56 genes in the double mutants (Figure 4A and Supplemental Table 3). Among the latter group, we found *daf-9,* encoding the cytochrome p450 that integrates inputs from TGF-β and insulin/IGF-II pathways to regulate DAF-12 activity during dauer development (49). We confirmed the increased expression of *daf-9* upon *chd-7* loss by RT-qPCR (Figure 4B). We hypothesized that *daf-9* expression levels must be critical in the decision to develop as either fertile adults or to enter diapause, and therefore the increased expression of *daf-9* in the *chd-7;daf-2* double mutants may be preventing full execution of the dauer program. Consistent with this hypothesis, we found that depletion of *daf-9* in *chd7*;*daf-7* animals restored the dauer arrest phenotype at the non-permissive temperature of 25°C (Figure 4C). However, these animals still appeared to be detergent-sensitive partial dauers (Figure 4D), since *daf-9* itself is required for proper dauer morphogenesis (39). Thus, we posit that the inability of *chd-7* mutants to fully repress *daf-9* may be sufficient to activate DAF-12 and inhibit dauer formation in both the TGF-β and IIS pathways (Figure 4E).

**Figure 4:**
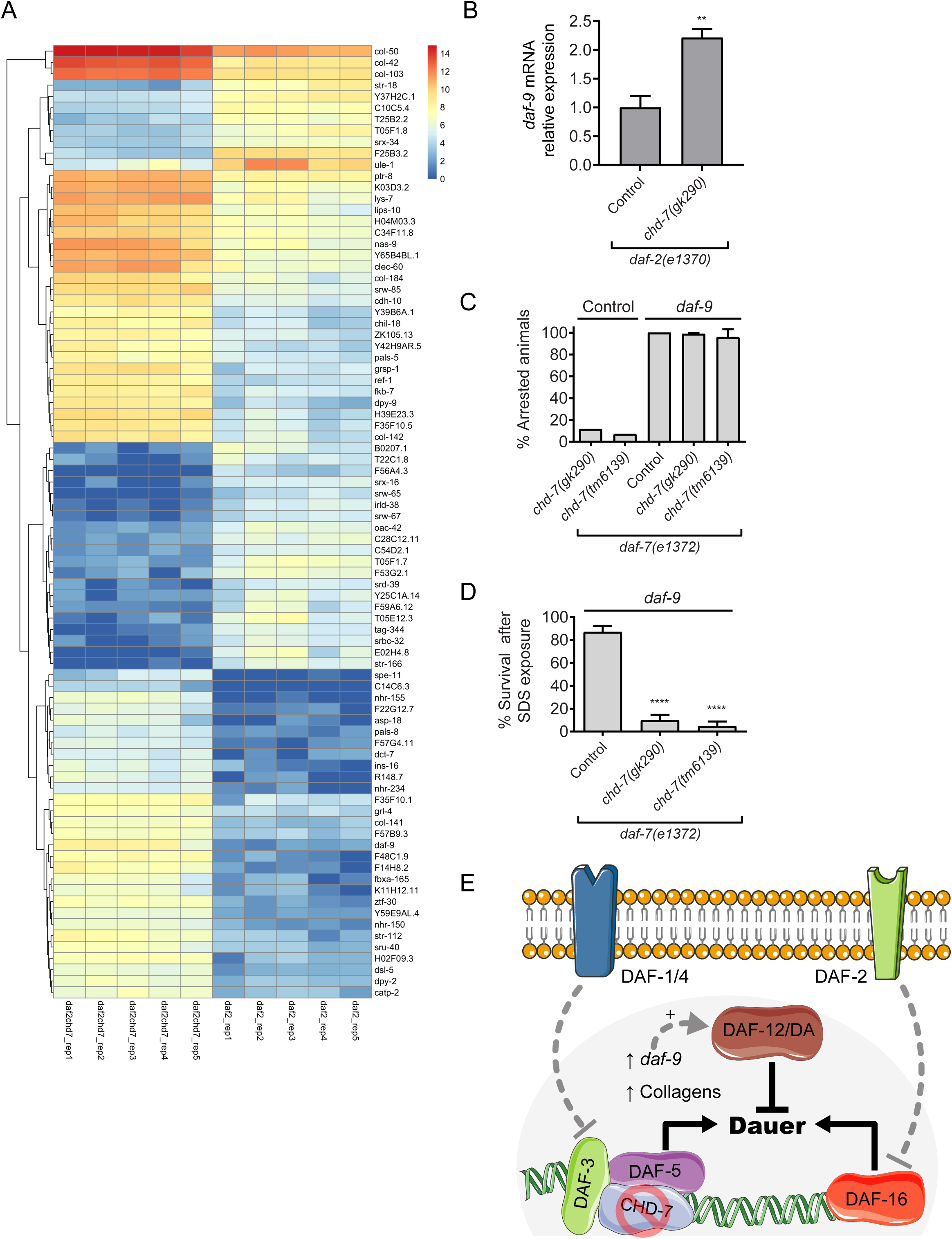
RNA-Seq analysis of transcriptome changes in *chd-7(gk290)* mutant dauers. A) Heat map of expression values for the 84 differentially expressed genes (DEGs, cutoff of 0.05 on FDR). DEGs were determined using DESeq2 (version 1.20.0). The color scale represents row z-score. Hierarchical clustering of the DEGs is represented by dendrograms at the left. B) Expression levels of *daf-9* mRNA are increased in *chd-7(gk290);daf-2(e1370)* partial dauers. Error bars indicate standard error from three repeats with different biological samples. Two-tailed unpaired t-test. **p <0.01. C) *daf-9* knockdown in *chd-7*;*daf-7(e1372)* animals rescues dauer arrest at 25°C. 2-3 young L4’s were plated on to freshly seeded plates with either *daf-9* or Control empty vector RNAi and allowed to lay eggs. After 72h, the adults were removed and the proportion of progeny that arrested as dauers was calculated. Each dot represents a plate of at least 34 animals (n>598 total animals/ strain). Bars and horizontal black lines represent mean percentage with SD. One-way ANOVA. D) *daf-9* RNAi rescues arrest in *chd-7*;*daf-7(e1372)* animals but leads to partial dauers. Arrested animals grown at 25°C on *daf-9* RNAi plates were treated with 1% SDS for 30 min and survival was scored. n>159 total animals/ strain Each dot represents a plate of at least 30 animals and a minimum of 159 total animals per strain tested. Bars and horizontal black lines represent mean percentage with SD. One-way ANOVA, ****p <0.0001, compared to *daf-7(e1372)*. E) Proposed mechanism of action for *chd-7* in dauer development. Under dauer-inducing conditions, CHD-7 forms a transcriptional complex with DAF-3/DAF-5 to repress *daf-9* expression. In *chd-7* mutants, *daf-9* expression prevents dauer formation driven by reduced activity of both DAF-7/TGF-β and DAF-2/IIS signaling pathways, presumably by binding dafachronic acids (DA) and subsequent activation of DAF-12.

The partial dauer larvae that arise from knockdown of *chd-*7 are short and thick and lack dauer alae, indicative of cuticle defects (Figures 1A and 1G). Thus, we reasoned that *chd-7* might also regulate target genes that are required for dauer morphogenesis. Further analysis of our transcriptomic data showed that 10% of the DEGs were collagens (*col-103*, *col-50*, *dpy-2*, *col-184*, *col-141*, *col-142*, *col-42* and *dpy-9*), which are structural components of the cuticle. All these collagens showed increased expression in the *chd-7*-mutant dauer larvae (Figure 4A). FPKM data from Modencode libraries indicates that each of these collagens show very low expression during dauer and are expressed during various stages of reproductive development, suggesting that repression of the genes must be important for dauer development (50). Thus, our results suggest that CHD-7 is required for repression of dauer-specific collagens. These results are consistent with growing evidence that the TGF-β pathway is a major regulator of collagen deposition (35, 51). In fact, it was recently reported that two of our target genes, *col-141* and *col-142*, contain SMAD-binding elements (SBEs) and their expression is directly regulated by the TGF-β pathway to determine body size (52).

In addition to the transcriptomic analysis of dauers, we analyzed CHD-7’s CHIP-seq data generated by the ModEncode project in young adults (Supplemental Table 4). Because the expression profiles of adult TGF-β and IIS mutants show significant overlap and co-regulation of many DAF-16 target genes (28), we compared this analysis with DAF-16 CHIP-seq data also from ModEncode in L4 larvae. To our surprise, we found that both transcriptional regulators share a significant number of genes (Supplemental Figure 5). Along with our observation that *chd-7* is necessary for lifespan extension of *daf-2* mutants, these data indicate that *chd-7* could modulate longevity by regulation of the IIS pathway, possibly through direct regulation of target genes.

### Chd7 regulates *col2a1* during *Xenopus* embryogenesis

Type-II collagen is an extracellular matrix (ECM) protein conserved in all multicellular animals, which forms fibrils (53) and has fundamental roles in development and tissues homeostasis (54). In vertebrates, the fibrillar type-II collagen is the major structural protein of cartilage and plays a prominent role in cranial development in multiple organisms (38, 55). Collagen, type-II, alpha 1 (Col2a1) is the major component of the cartilage matrix having a structural function and being an important extracellular signaling molecule for regulation of chondrocyte proliferation, metabolism, and differentiation (56–58). The African frog *Xenopus laevis* is a well-established model to study vertebrate facial disorders which often arise from defects in neural crest development and migration (37). In *Xenopus* embryos, prior studies established that Chd7 regulates neural crest specification and migration and its depletion recapitulates the craniofacial defects seen in CHARGE patients (36). To investigate whether Chd7 has a conserved mechanism of action and regulates collagen expression in vertebrates, we used a previously validated morpholino to induce *Xenopus* Chd7 knockdown (*chd7*-MO) (36). In *Xenopus,* Col2a1 is essential for normal development of the skeleton and its expression is restricted to the cartilaginous skeleton of the tadpole and adult frog (59). In *chd7*-MO injected embryos, RT-qPCR revealed a significant reduction of *col2a1* mRNAs as compared to control-injected animals (Figure 5A).

**Figure 5:**
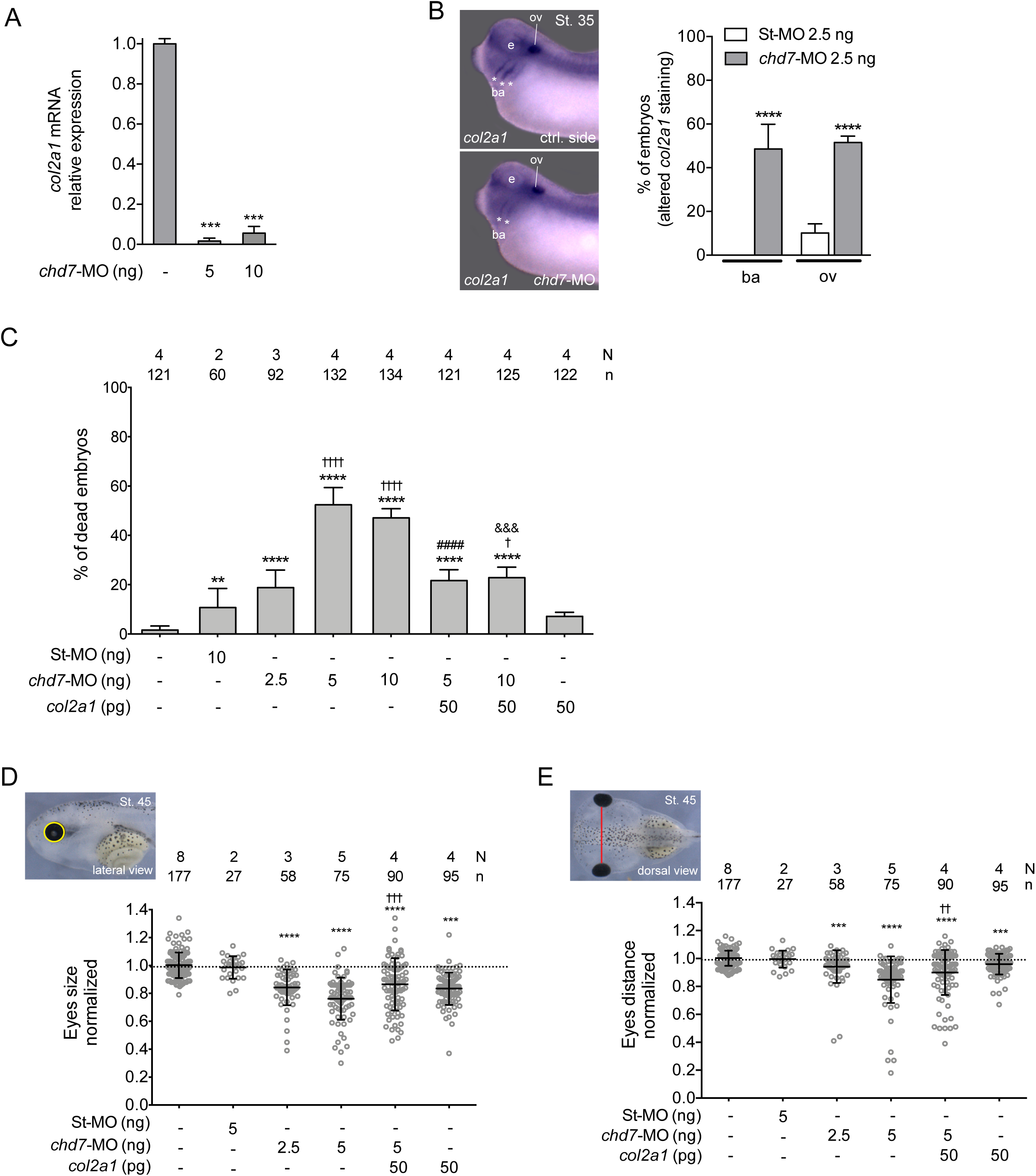
Expression of *col2a1* rescues Chd-7 knock-down in *Xenopus* embryos. A) Expression levels of *col2a1* mRNA are reduced in *chd7*-MO injected embryos. Both blastomeres of 2-cell staged embryos were injected with 5 or 10 ng of *chd7*-MO. Error bars indicate standard error from two repeats of the PCR reaction with different biological samples. One-tailed paired t-test, \*\*\**p*< 0.001. B) *Col2a1* expression domain is altered in the branchyal arches (ba) and otic vesicle (ov) of Chd7-depleted embryos. *In situ* hybridization of stage (St.) 35 embryos for *col2a1*. Lateral views, anterior to the left. One D1 blastomeres of 8-cell staged embryos was injected with 2.5 ng of St-MO or *chd7*-MO (N = 2; 42 and N = 3; 71, respectively) (N = number of experiments; number of embryos). Top and bottom panels are control and injected sides of the same representative *chd7*-MO injected embryo, respectively. e: eye. The graph is a quantification of the results. Reduced *col2a1* staining was observed in 18% of St-MO and 77% of *chd7*-MO injected embryos. Data on graph is presented as means with standard error. Fisher’s exact test (\*\*\*\**p*< 0.0001). * represents comparison to St-MO group. C) Lethality in Chd-7-depleted embryos is rescued by over-expression of *col2a1*. Graph showing the percentage of dead embryos. Both D1 blastomeres of 8-cell staged embryos were injected as indicated in the graph and scored for survival at St. 45. Data on graph is presented as means with standard error. Fisher’s exact test (^†^*p*< 0.05, \*\**p*< 0.01, ^&&&^*p*< 0.001, ****^,††††,####^ *p*< 0.0001). * represents comparison to uninjected group, ^†^ represents comparison to St-MO group, ^#,&^ represent comparison to *chd7*-MO 5 ng and 10 ng, respectively. D-E) Craniofacial morphometric analysis of *Xenopus* tadpoles at St. 45. Embryos were injected as indicated in panel C. Each dot represents a single embryo. Means and standard deviation are indicated. One-way ANOVA and Tukey’s multiple comparisons test. D) Quantification of the eye size (top left panel). ***^,†††^*p*< 0.001, \*\*\*\**p*< 0.0001. E) Quantification of the eye distance (top left panel). ^††^*p*< 0.01, \*\*\**p*< 0.001, \*\*\*\**p*< 0.0001. Lateral views, anterior to the left. * represents comparison to uninjected group and ^†^ represents comparison to *chd7*-MO 5 ng injected group. N = number of experiments, n = number of embryos.

For targeted disruption of Chd7 function, we injected the dorsal-animal (D1) blastomeres of 8-cell stage embryos fated to contribute to the dorsal anterior structures. *In situ* hybridization of unilaterally injected embryos showed alterations in *col2a1* expression in the branchial arches and/or the otic vesicle (ear vesicle) (59) upon *Chd7* depletion relative to embryos injected with a control morpholino (St-MO) (Figure 5B). The injection of *chd7*-MO into both D1 blastomeres of 8-cell stage embryos with doses between 1.25 ng and 2.5 ng showed 10-20% embryonic mortality (Figure 5C and data not shown). The higher doses of 5 ng and 10 ng led to >50% lethality (Figure 5C). We next sought to investigate if the mortality was related to collagen deficits and therefore conducted rescue experiments by co-expressing *Xenopus col2a1* mRNA with the morpholino. As shown in Figure 5C, coinjection of *col2a1* mRNA substantially improved (∼50%) embryo survival relative to the injection of *chd7*-MO alone. We then asked if ectopic *col2a1* mRNAs could suppress the craniofacial defects associated with *chd7* loss-of-function. Initially, we analyzed the gross morphology of the surviving stage 45 tadpoles and observed a high incidence of craniofacial malformations (83%) in Chd7-depleted animals (Supplemental Figure 6A and 6C) and a significant reduction upon *col2a1* mRNA coinjection (43%) (Supplemental Figure 6B). Next, we examined eye size and eye distance, since microthalmia and midline defects are often associated with CHARGE syndrome (60) and are recapitulated in *Xenopus* embryos (36). Both eye size and distance between eyes were reduced in Chd7-depleted tadpoles and were partially rescued by *col2a1* mRNA expression (Figure 5D and 5E). Therefore, expression of *col2a1* ameliorated the phenotypes associated with pathogenic Chd7 suggesting that collagen is a conserved and important target of this protein.

## DISCUSSION

We initially identified *chd-*7 in chromatin immunoprecipitation (ChIP)-chip studies of the nuclear receptor DAF-12 (34), a transcription factor regulating worm aging, development, and dauer formation (61), and found it to be critical for *daf-2* mediated dauer development. Comparison of the *chd-7* alleles *gk290* and *gk306* showed a critical role for the BRK domain in dauer development and longevity (Figures 1F-H, 2A and 2C). In CHARGE patients, deletions or mutations within the BRK domains of CHD7 are sufficient to elicit all the features characteristic of the disease, underscoring the importance of this domain (62). The phenotypic differences between the worm alleles highlight the potential of the worm to delimit functional domains of CHD-7 that contribute to disease pathology. In mice, homozygous mutations in *Chd7* lead to embryonic lethality at E10.5, in part because *Chd7* is necessary for early brain development (36). In worms, the presumptive null allele c*hd-7(tm6139)* and *scd-3(sa253)* were viable but showed reproductive defects, such as improper gonad proliferation and migration, reduced fecundity, male tail defects and hermaphrodite vulval defects (31), indicating that *C. elegans* are more able to tolerate loss of CHD-7 than mice or humans. Thus, our studies establish *C. elegans* as an animal model to study the mechanisms underlying the developmental defects observed in pathogenic Chd7.

Both a *chd-7* mutation and *chd-7* overexpression shortened the lifespan of otherwise wild-type worms, suggesting that CHD-7 proteins levels must be tightly regulated to ensure proper development. Of note, Chd7 is the most commonly amplified gene in tumors amongst the CHD superfamiliy members, and its overexpression is associated with aggressive subtypes of breast cancer and poor prognosis (63). Thus, in humans, as well as worms, gain-of-function phenotypes are associated with Chd7/*chd-7* overexpression.

Our genetic epistasis analyses placed CHD-7 downstream of the TGF-β-like DAF-7, the type I receptor DAF-1, and the R-SMAD DAF-14 placing CHD-7 at the level of the Co-Smad DAF-3 and the Sno/Ski repressor DAF-5, which are also Daf-d (64). While *daf-3* and *daf-5* completely suppress dauer formation in a *daf-7* background at 25°C, *daf-5* fails to suppress dauer formation in TGF-β mutants at 27°C (26), supporting a role for CHD-7 at the DAF-3/DAF-5 step in the pathway (Figures 3A and 3B). Interestingly, these transcriptional regulators have opposite effects on *daf-2*-induced longevity: *daf-3* enhances *daf-2(e1370)* longevity while *daf-5* mutations suppresses it (30). We observed that *chd-7* suppresses *daf-2* longevity (Figure 2C), like *daf-5(e1386)*. Together these data suggest that CHD-7 either regulates *daf-5* expression or directly interacts with DAF-5 to regulate downstream target genes, or both. In mice, CHD7 was shown to physically interact with SMAD1 and form a transcriptional complex with SMAD4, the mammalian ortholog of DAF-3 (65). We therefore speculate that DAF-3, DAF-5, and CHD-7 may be in ternary complex that regulates *daf-9* expression for dauer entry (Figure 4C). Interactome mapping of the TGF-β pathway also identified SWSN-1, a SWI/SNF subunit component of the BAF complex, as a physical interactor of DAF-3 (66). CHD7 interacts with human and *Xenopus* PBAF (polybromo- and BRG1-associated factor-containing complex) to control neural crest gene expression (36). In worms, both *swsn-1* and *chd-7* fail to develop normal dauers in *daf-2* and *daf-7* mutants (Supplemental Figure 3 and (67)). Thus, we envision that CHD-7 may work together with the BAF complex and DAF-3/DAF-5 to control gene expression of target genes critical for dauer formation.

DAF-9 activity is necessary and sufficient for the decision to enter diapause: reduced activity of TGF-β and IIS pathways leads to *daf-9* repression and dauer development. Conversely, *daf-9* expression in the hypodermis is sufficient to inhibit diapause, driving reproductive programs in *daf-7* mutants (29, 32). Ectopic *daf-9* expression also drives reproductive programs in the weak *daf-2(e1368)* allele, but only partially suppresses *daf-2(e1370)* diapause leading to arrest as L3 or early L4 larvae (32). Based on these observations, we speculate that *daf-9* misexpression explains a subset of the *chd-7* phenotypes observed herein, including the partial dauer phenotype, gonad migration defects, and vulval protrusions, all of which overlap with published *daf-9* phenotypes (32, 33, 68). Interestingly, *daf-9* is both upstream and downstream of *daf-12* for dauer formation (*32, 33*). Our data demonstrated that DAF-12 regulates *chd-7* expression and CHD-7 in turn regulates *daf-9* expression. Therefore, we propose that CHD-7 belongs to the transcriptional complex that regulates the feedback loop between *daf-12* and *daf-9*.

Among the differentially expressed genes, we noted the presence of several G protein coupled receptors (GPCRs), which could be candidate chemoreceptors for dauer pheromone or environmental cues and thus may be required for either dauer entry or maintenance (42). Consistent with our transcriptomic studies, *Liu et al.* analyzed the expression differences between L2/L3 larvae and TGF-β mutants undergoing diapause and found enrichment of collagen genes, GPCRs and *daf-9*, further validating our placement of *CHD-7* in the TGF-β dauer signaling pathway (29). In a more recent paper, also was established that GPCR gene expression increases significantly during dauer commitment (69).

In *C. elegans*, the TGF-β family pathway has diversified and has unique ligands and effectors for the control of dauer induction and aspects of somatic development. Mutations in the TGF-β ligand *dbl-1* and its downstream components cause a small body size (Sma phenotype) (47, 48). Consistent with *chd-7* being a component of the TGF-β signaling pathway, *chd-7(gk290)* and *chd-7(tm6139)* are significantly shorter than wild-type worms (Figure 3C). This phenotype is also observed in *scd-3* mutants (31). The *dbl-1* pathway mutants have defects in male tale development (48), a phenotype shared in *scd-3* (31) and *chd-7* mutant animals (data not shown and (31)). Therefore, our results suggest that CHD-7 acts as a regulator of both the *daf-7* and *dbl-1* branches of the TGF-β pathway.

Craniofacial anomalies seen in CHARGE patients involve tissues deriving from cells of the neural crest, including craniofacial cartilage and bone, heart, ears and eyes (60, 70). *Xenopus laevis* is an established model to understand fundamental questions of craniofacial biology (37). *Chd7* knockdown in frogs and in zebrafish (11, 36), leads to defects in neural crest specification and migration, which was recently been recapitulated in human embryonic stem cells (hESCs) (71). Our RNA-seq analysis revealed that CHD-7 regulates expression of genes from the cuticle, a modified extracellular matrix (72). Here, we show a role for Chd7 in regulating *col2a1* expression, a type II collagen and the major component of cartilage (55). Interestingly, regulation of *col2a1* by Chd7 is also observed in zebrafish (73). While additional studies are required, we speculate that Chd7 could regulate *col2a1* in a complex with the transcription factor Sox10 (74, 75) or through the TGF-β signaling pathway (35, 76). Supporting the latter mechanism, it was demonstrated in chondrocytes that TGF-β regulates *col2a1* expression (77, 78). Taken together, our data supports a model in which collagen misexpression by pathogenic Chd7 leads to craniofacial defects and embryonic lethality.

## MATERIALS and METHODS

### *C. elegans* strains

All strains were grown and maintained on standard nematode growth medium (NGM). In Argentina (Hochbaum lab), plates were supplemented with 0.1 mg/ml streptomycin and 100 U/ml Nystatin using the *E. coli* OP50-1 strain as the food source. In Pittsburgh (Yanowitz lab), strains were grown on standard NGM seeded with OP50. Temperature-sensitive strains were maintained at 16°C and shifted to the non-permissive temperature of 25°C to induce dauer formation, unless otherwise indicated. All other strains were maintained at 20°C. All *daf-7* strains were maintained at low density to prevent dauer induction at 16°C. Dauer studies were performed in both the Yanowitz and Hochbaum laboratories to validate results. All *chd-7* alleles were outcrossed at least 5 times prior to use.

Strains utilized in this study are listed in Supplemental Table 1. Standard genetic crosses were used to make double or triple mutants. The presence of mutant alleles was confirmed a) by the *daf-c* phenotypes in animals heterozygous for additional mutations and b) by PCR and/or sequencing for all additional mutations.

### RNAi screen for dauer suppressors

All RNAi clones were picked from the Ahringer bacterial feeding library (79). These *E. coli* clones were seeded on NGM plates supplemented with 1 mM of IPTG (Isopropyl β-D-1-thiogalactopyranoside) and 0.1 µg/ml ampicillin, and used for inducing RNAi by the feeding method (80).

GL228 [*rrf-3(pk1426)*] II;*daf-2(e1371)* III] eggs were placed in 24-well RNAi plates seeded with bacteria expressing the dsRNA of interest. Worms were maintained for 5 days at 15°C until adulthood, then were transferred to an identical 24-well plate to lay eggs for 5 h. Adults were removed and the eggs were incubated at 25°C for 4 days to allow formation of dauers. *daf-16(RNAi)* and the empty vector were used as controls. Proper dauer formation was assessed by observation in a dissecting microscope and by 1% SDS resistance. RNAi clones that caused abnormal dauer phenotypes were validated in *daf-2(e1370)* worms. Identity of the dsRNA was confirmed by sequencing (Macrogen, Korea).

### *daf-9* suppression assays

For preparation of *daf-9*(RNAi) plates, bacterial cultures were grown overnight in LB with 10 µg/ml tetracycline and 50 µg/ml carbenicillin and induced with 4 mM IPTG for 4 h. Cultures were spun down, suspended in 1/10 volume and 30 µl of bacteria were seeded on 3 cm NGM plates made with 1 mM IPTG and 50 µg/ml carbenicillin. Plates were grown overnight at 25°C and stored at 4°C for no more than 2 weeks prior to use.

L4 stage animals were placed on 3 cm RNAi plates. Two worms per plate were used for *daf-7(e1372)*, while three worms were used for *daf-7;chd-7(gk290)* and *daf-7;chd-7(tm6139)*. After 72 h, the adults were removed and plates were replaced at 25°C for 2-3 days. The total number of dauers, L4s, and adults were then assessed.

### SDS survival assay

Young adults were transferred to seeded plates and permitted to lay eggs for 5 days at 25°C. The arrested progeny were then washed off plates with M9 (22 mM KH_2_PO_4_, 42 mM Na_2_HPO_4_, 85.5 mM NaCl, 1 mM MgSO_4_) into 15 ml glass conical tubes. Collected animals were washed 2-3 times with M9 and the excess liquid was aspirated off. Animals were then treated with 2 ml of 1% SDS for 30 min on a nutator at 25°C. Following incubation, the samples were washed 3 times with M9 and any excess liquid was aspirated off. Animals were aliquoted to 5 seeded plates with 50-70 worms per plate and allowed to recover at 16°C. The recovered animals were then quantified, and the percentage recovered was calculated. This was repeated 3 times for each strain tested. Comparisons were performed using the χ-squared Test in R; P values were corrected using the Bonferroni method for multiple comparisons.

### *fog-2* mating assay

To determine if males were capable of siring offspring, 8 males from each strain tested were plated with 4 *fog-2(q71)* adult females on seeded 10 cm plates. After 24 h, *fog-2* females were transferred to new plates and within 48 h the proportion of fertile females were scored. Assay was repeated two times.

### Lifespan assays

All lifespan experiments were conducted by transferring 1 day-old adults from 15°C to 20°C for the remainder of the lifespan assay. NGM plates were seeded with *E. coli* OP50-1. ∼150 L4 hermaphrodites were transferred to 5 plates per experiment. Every 48 h, animals were scored as alive, dead or censored (animals that exploded, died from bagging or dried out at the edges of the plates). Animals were considered dead when they did not respond to a soft touch to the head with a pick. To prevent the progeny from interfering with the assay, adults were transferred to fresh plates every 48 h until egg production ceased. For *glp-1(e2144)* assays, eggs were kept at 20°C for 4 h and then transferred at 25.5°C for 72 h to induce sterility and switched to 20°C for the remainder of the experiment. Lifespan data were analyzed using the Kaplan-Meier method. Statistics were calculated using the Mantel-Cox nonparametric log-rank method using OASIS2 (81).

### Pathogen resistance assay

The pathogenic bacterial strain used in this study was *Pseudomonas aeruginosa* (strain PA14). This strain was streaked from a frozen stock onto an LB agar plate, incubated at 37°C overnight and then kept at 4°C (shelf-life of one week). For survival assay, a single PA14 colony was inoculated in King’s broth and incubated at 37°C overnight with shaking. 20 µl of this culture was seeded onto slow killing NGM plates (containing 0.35% peptone instead of 0.25%) and incubated for 24 h at 37°C. The plates were then left to sit at room temperature for 24 h. The following day, 150 L4 hermaphrodites grown at 15°C were distributed onto five PA14 plates and incubated at 25°C. Survival was monitored at intervals of 6-12 h and live, dead and censored animals were recorded. Data and statistics were analyzed using the Kaplan-Meier method as described in the section “Lifespan assays”.

### Microscopy and fluorescence imaging

For imaging dauers and adults, worms were immobilized in a 25 mM sodium azide solution on fresh 4% agarose pads and imaged at 10x and 20x magnification. Images were collected using a Zeiss Axioplan Imaging Microscope with a DIC system and Zeiss Plan-Neofluar 10x and 20x objectives lens. Images were acquired with a Micropublisher 3.3 camera (Q Imaging). ImageJ (NIH) software was used to quantify worm size.

For imaging *pCHD-7::mCherry* fluorescence, worms were immobilized with 1 mM levamisole on fresh 2% agarose pads and imaged immediately using a Nikon A1r confocal microscope equipped with a 40x PLAN APO oil objective. ImageJ (NIH) software was used to quantify fluorescence intensity.

For imaging gonads in *daf-2(e1370)* and mutants, young adults were transferred to seeded plates and permitted to lay eggs for 5 days at 25°C to form dauers, then collected and fixed with Carnoy’s solution (75 µl EtOH, 37.5 µl Acetate, 12.5 µl Chloroform) and stained with 5 mg/ml DAPI (4′,6-diamidino-2-phenylindole) in PBS. Animals were imaged as 0.5 µm Z-stacks with the Nikon A1r Confocal Microscope with 40x and 60x plan APO oil objectives.

For imaging *ajm-1::GFP*, young adults were transferred to seeded plates and allowed to lay eggs for 5 days at 25°C to form dauers. The arrested progeny was then washed off plates with M9 into 15 ml glass conical tubes. Collected animals were washed 2-3 times with M9 and the excess liquid was aspirated off. Animals were then immobilized with levamisole and the seam cells were imaged as Z-stacks by confocal microscopy as described immediately above.

For scanning electron microscopy, worms were fixed by immersing in 2.5% glutaraldehyde in PBS for several hours. Worms were washed 3X in PBS then post fixed in aqueous OsO_4_ for 1h, then washed 3x in PBS, dehydrated through a graded series of ethanols (30-100%) then chemically dried with 2x 10 min incubations in hexamethyldisilazane. Dried worms were sprinkled onto copper double stick tape on aluminum stubs, sputter coated with 3.5 nm gold-palladium alloy then evaluated on a JEOL JEM 6335F microscope at 5 kV.

### Library preparation and RNA-seq

*daf-2(e1370)* and *chd-7(gk290);daf-2(e1370)*, synchronized eggs were kept at 25°C for 10 days and resulting dauers were collected and frozen. Total RNA was extracted with TRIzol (Invitrogen) following the kit’s protocol. cDNA library was prepared with NEBNext Ultra II RNA library prep kit for Illumina (New England Biolabs), and the sequencing carried out using Illumina’s HiSeq-2500 sequencer with single-end mode and read length of 50 bp. Five replicates for *daf-2(e1370)* vs. *chd-7(gk290);daf-2(e1370)* were sequenced. For data assessment, a quality control with FastQC software (version 0.11.5) was used. First the raw reads that aligned against the *E. coli* genome (K12 genome) were removed. The remaining sequences were aligned against the reference genome of *C. elegans* WS260 using STAR (version 2.5.4a). The number of mapped reads to genes was counted using Htseq (version 0.9.1). Finally, the DEGs were determined using DESeq2 (version 1.20.0) with a cutoff of 0.05 on False Discovery Rate (FDR). R version 3.5.0 (2018-04-23) and Bioconductor version 3.7 with BiocInstaller version 1.30.0 were used. Heatmaps were generated using pheatmap package (version 1.0.12) with hierarchical clustering on the rows with the default options.

### Genomic Sequencing of *scd-3*

A 6 cm plate replete with *scd-3* gravid animals was washed with 1 ml M9 and collected in a glass conical tube. Worms were washed extensively and then placed on a nutator for 1 h to allow remaining gut bacteria to be passed into the medium. Worms were again washed 3-4 times and settled by gravity. The worm pellet was transferred to 1.5 ml tubes and genomic DNA was prepared according to the manufacturer’s protocol with the Purelink Genomic DNA Kit (Invitrogen) except that after addition of digestion buffer, worms were pulverized with a microfuge hand dounce prior to incubation at 55°C. Sequencing was performed by Psomagen, Qiagen CLC Genomics Workbench was used to align the DNA against WBcel235 and view the variant.

### Oil-Red-O (ORO) staining

Dauers grown at 25°C for 5 days were stained for lipids using ORO, as previously described (82). Animals were mounted on a 4% agarose pad and observed in a Zeiss Axioplan brightfield microscope equipped with a Micropublisher 3.3 camera (Q Imaging). Image J (NIH) was used to quantify the amount of lipids in each animal. At least 20 animals of each strain were quantified. Statistical differences was determined by Student’s t-test.

### L1 survival

Experiments were done at 20°C. Eggs were obtained by bleaching of gravid adults and kept under gentle shaking in 5ml sterile M9 supplemented with 0.1µg/ml streptomycin to hatch (16h). To normalize population density, resulting L1 larvae were diluted to obtain 1 larva/µl in 5ml of M9 and kept under constant agitation for the remainder of the experiment. Every 48h, a 100µl aliquot of L1 larvae were spotted onto a NGM plate and then incubated for 72h. Animals were scored as alive if developed beyond the L2 stage. Percentage of the population alive was normalized to day 1 seeded L1 larvae.

### Gonad staining

Late L4/Young Adult animals from the relevant strains were collected and fixed with Carnoy’s solution (75 µl EtOH, 37.5 µl Acetate, 12.5 µl Chloroform) and stained with 5 mg/ml DAPI (4′,6-diamidino-2-phenylindole) in PBS. Animals were imaged as 0.5 µm Z-stacks with the Nikon A1r Confocal Microscope with 20x objective.

### ChIP-seq analysis

Data from CHD-7 and DAF-16 ChIP-seq generated by ModEncode project was analyzed. Reference for CHD-7-eGFP: https://www.encodeproject.org/experiments/ENCSR010MNU/. Reference for DAF-16-eGFP: https://www.encodeproject.org/experiments/ENCSR946AUI/. Peaks were downloaded in ce10 and annotated using Homer software. Gene lists from the peak calling were generated and used to compare CHD-7 and DAF-16.

### *Xenopus laevis* embryo manipulation and microinjections

*Xenopus* embryos were obtained by natural mating. Adult frogs’ reproductive behavior was induced by injection of human chorionic gonadotropin hormone. Eggs were collected, de-jellied in 3% cysteine (pH 8.0), maintained in 0.1 X Marc’s Modified Ringer’s (MMR) solution and staged according to Nieuwkoop and Faber (83). The embryos were placed in 3% ficoll prepared in 1 X MMR for microinjection. Chd7 morpholino (*chd7*-MO: 5′-AACTCATCATGCCAGGGTCTGCCAT-3′) specificity has been previously characterized (36). *Chd7*-MO and Standard Control morpholino (St-MO) were provided by Gene Tools, LLC. The cDNA of *X. laevis col2a1* was amplified by PCR from pCMV-Sport 6-*col2a1* (Dharmacon) with primers M13F and M13R. The PCR fragment was digested with *EcoR*V and *Not*I and cloned into pCS2+ previously digested with *Stu*I and *Not*I. Capped mRNAs for *col2a1* were transcribed *in vitro* with SP6 using the mMessage mMachine kit (Ambion) following linearization with *Not*I. *Chd7*-MO and *col2a1* mRNA were injected into both D1 blastomeres of 8-cell staged embryos (82) for lethality and morphometrics analysis. *Chd7*-MO was injected into one D1 blastomeres of 8-cell staged embryos for analysis of *col2a1* expression.

Whole-mount *in situ* hybridization was carried out as previously described (85). pCMV-Sport 6-*col2a1* (Dharmacon) construct was linearized with *Sal*I and transcribed with T7 for antisense probe synthesis. Morphometrics analyses were done on fixed tadpoles using ImageJ (NIH) software. Morphometric measurements were normalized to the mean of the uninjected group in order to compare between independent experiments. For cartilage staining, stage 45 tadpoles were fixed with MEMFA for 24 hours at 4°C, dehydrated into 100% ethanol and stained in 0.01% Alcian blue 8GX in 70% ethanol/30% glacial acetic acid for three nights. Distaining was done in 100% ethanol followed by rehydration in 2% KOH. Animals were cleared in graded glycerol in 2% KOH and skulls were dissected under stereoscope. Images of whole embryos and skulls were collected with a Leica DFC420 camera attached to a Leica L2 stereoscope.

### RNA preparation and RT-qPCR

For *Xenopus* RNA extraction, 2-cell staged embryos were injected into both blastomeres with 5 ng or 10 ng of *chd7*-MO, collected at stage 23 and snap-frozen for later processing. Six embryos were combined for each treatment. For worm RNA extraction, dauers and partial dauers were developed for a week at 25°C. For all the samples, RNA was isolated using RNAzol (Molecular Research). RNA concentration of each sample was measured using a Nanodrop spectrophotometer and 250 ng of RNA was reverse transcribed using qScript cDNA SuperMix (QuantaBio). Quantitative PCR (qPCR) was preformed using the Forget-Me-Not qPCR Master Mix (Biotium) with a BioRad iCycler thermocycler. Amplification was performed using oligonucleotides designed with PerlPrimer software for *col2a1* (Forward 5’ TCCCTGTTGATGTTGAAGCC 3’; Reverse 5’CAATAGTCACCGCTCTTCCA 3’ and ODC primers have been previously described (86) (Forward 5’CAAAGCTTGTTCTACGCATAGCA 3’; Reverse 5’GGTGGCACCAAATTTCACACT 3’). The relative expression of *col2a1* was normalized to *ODC* expression and determined via the method previously described (87). *daf-9* was amplified with specific oligos (Forward 5’ATTCCCCACAAAACAATCGAAGAAT 3’; Reverse 5’ GAGATTCAAACACGTTTGGATCG 3’) and expression was normalized to the housekeeping gene *cdc-42* (Forward 5’CTGCTGGACAGGAAGATTACG 3’; Reverse 5’ CTGGGACATTCTCGAATGAAG 3’).

### Ethics Statement

*Xenopus laevis* experiments were carried out in strict accordance with the recommendations in the Guide for the Care and Use of Laboratory Animals of the NIH and also the ARRIVE guidelines. The animal care protocol was approved by the Comisión Institucional para el Cuidado y Uso de Animales de Laboratorio (CICUAL) of the School of Applied and Natural Sciences, University of Buenos Aires, Argentina (Protocol #64).

## Supporting information

ChIP-seq analysis

DEGs

Strains

Longevity

## ACKNOWLEDGMENTS

The authors would like to thank Bruno Moretti, Hernan Grecco, Mario Rossi, and Julie Kocherzat for experimental support. DMJ and ASC were supported by the Consejo Nacional de Investigaciones Científicas y Técnicas (CONICET) Doctoral Fellowship Program. DH laboratory was supported by the Agencia Nacional de Promoción Científica y Tecnológica of Argentina (PICT-2016-0269) and CONICET (PIP 1122015 0100731 CO). MCC laboratory was supported by the Agencia Nacional de Promoción Científica y Tecnológica of Argentina (PICT-2013-0381). Funding for this work was also provided by grants from the CHARGE Syndrome Foundation (JLY and DH), The Company of Biologists (DMJ), NIGMS (R01GM104007 to JLY), and NIA (R01AG051659 to AG). This work was supported in part by the Intramural Research Program of the NIH and the National Institute of Diabetes and Digestive and Kidney Diseases (SY).

## SUPPLEMENTAL FIGURES

**Figure S1:**
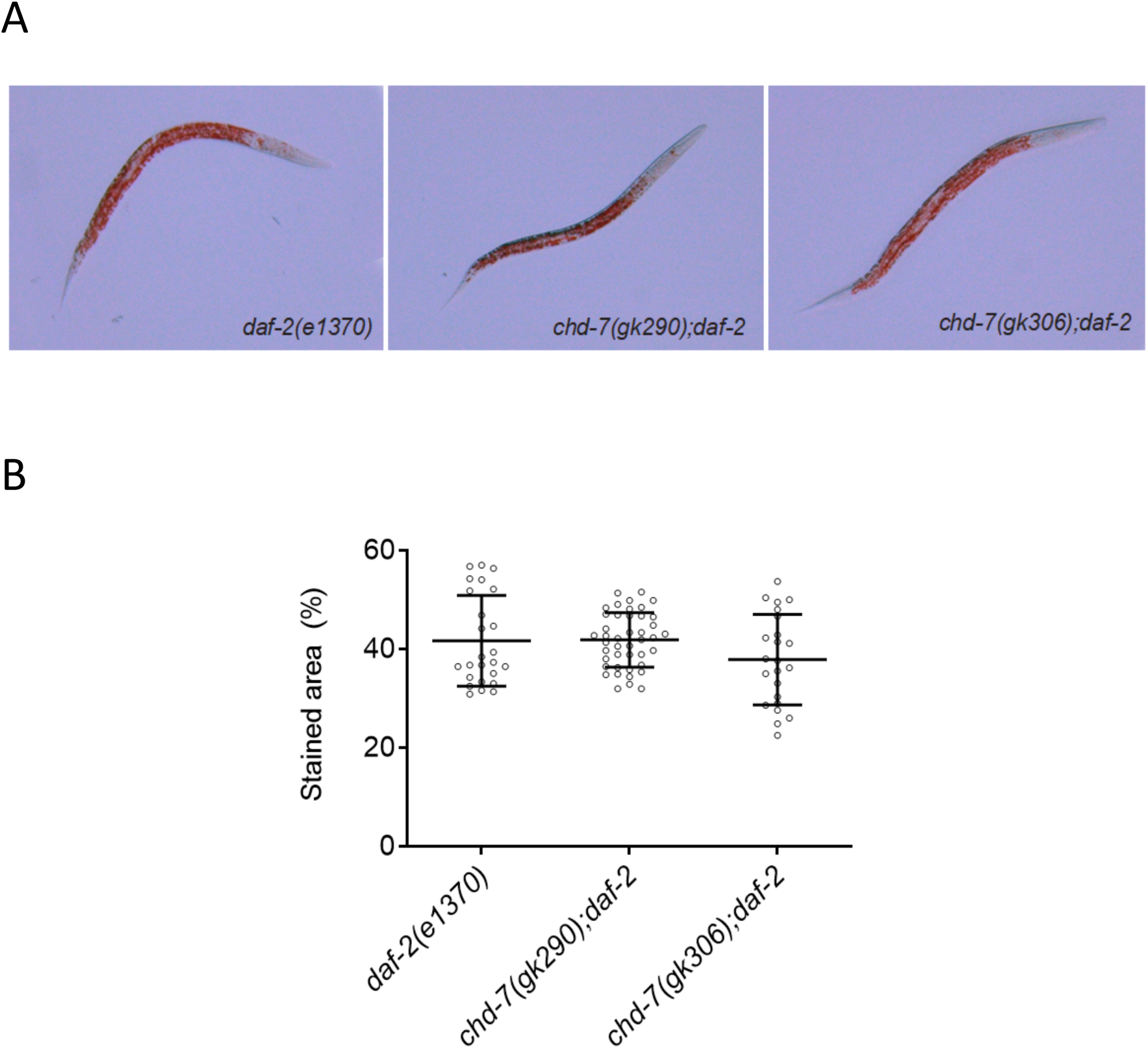
*chd-7* does not affects fat storage. Dauers of the shown genotypes were grown at 25°C for 5 days prior to lipid staining with Oil Red O (ORO). A. Representative photomicrographs of ORO-strained worms. B. Quantification of total area of ORO staining/ worm (Image J, see methods) reveals no significant differences in fat accumulation between control and *chd-7* mutants. Three biological replicates were scored with at least 16 individual dauers per replicate. Horizontal black lines represent mean with SD. P>0.05, Student’s t test.

**Figure S2:**
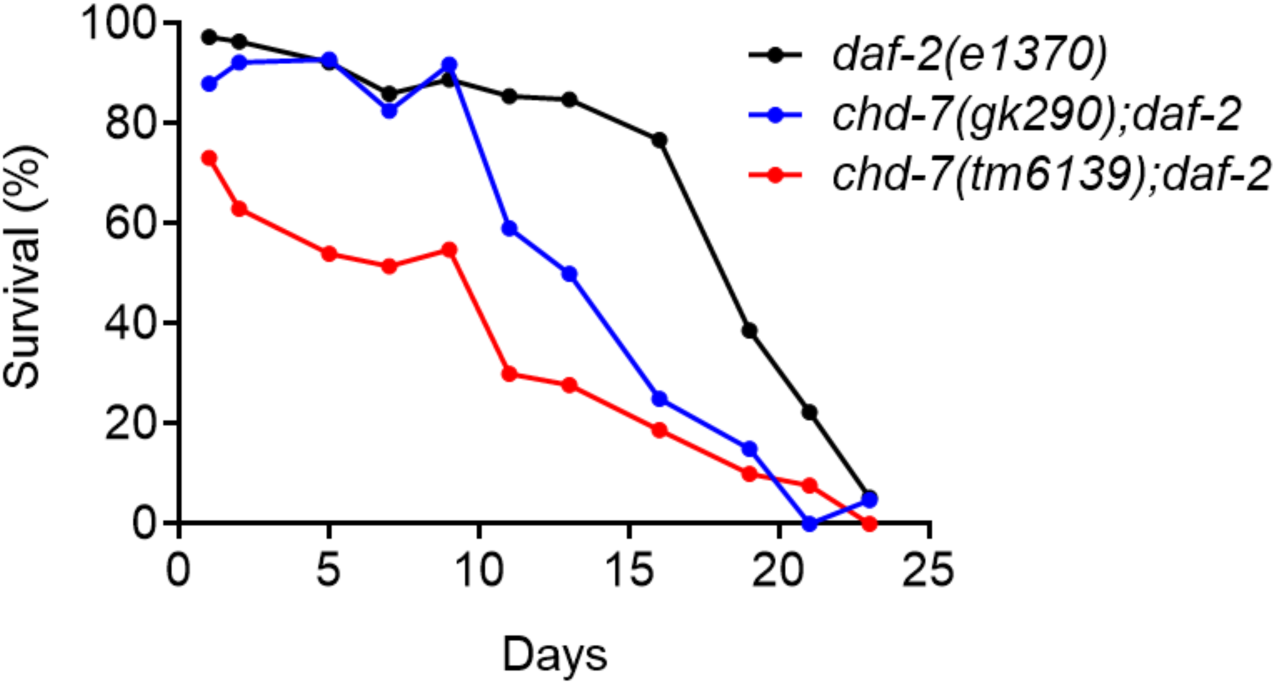
Loss of *chd-7* reduces L1 starvation survival in *daf-2(e1370)* mutants. L1 animals hatched from bleached egg preps into sterile M9 with 0.1µg/ml of streptomycin were diluted to a density of 1 larva/µl and kept at 20°C with constant agitation. Every 48h, 100µl aliquot was spotted onto a seeded NGM plate, and scored 72h later for development beyond the L2 larval stage. Percentage of the population alive was obtained by comparing the initial number of worms at t=0. Each dot represents at least 50 L1 worms.

**Figure S3:**
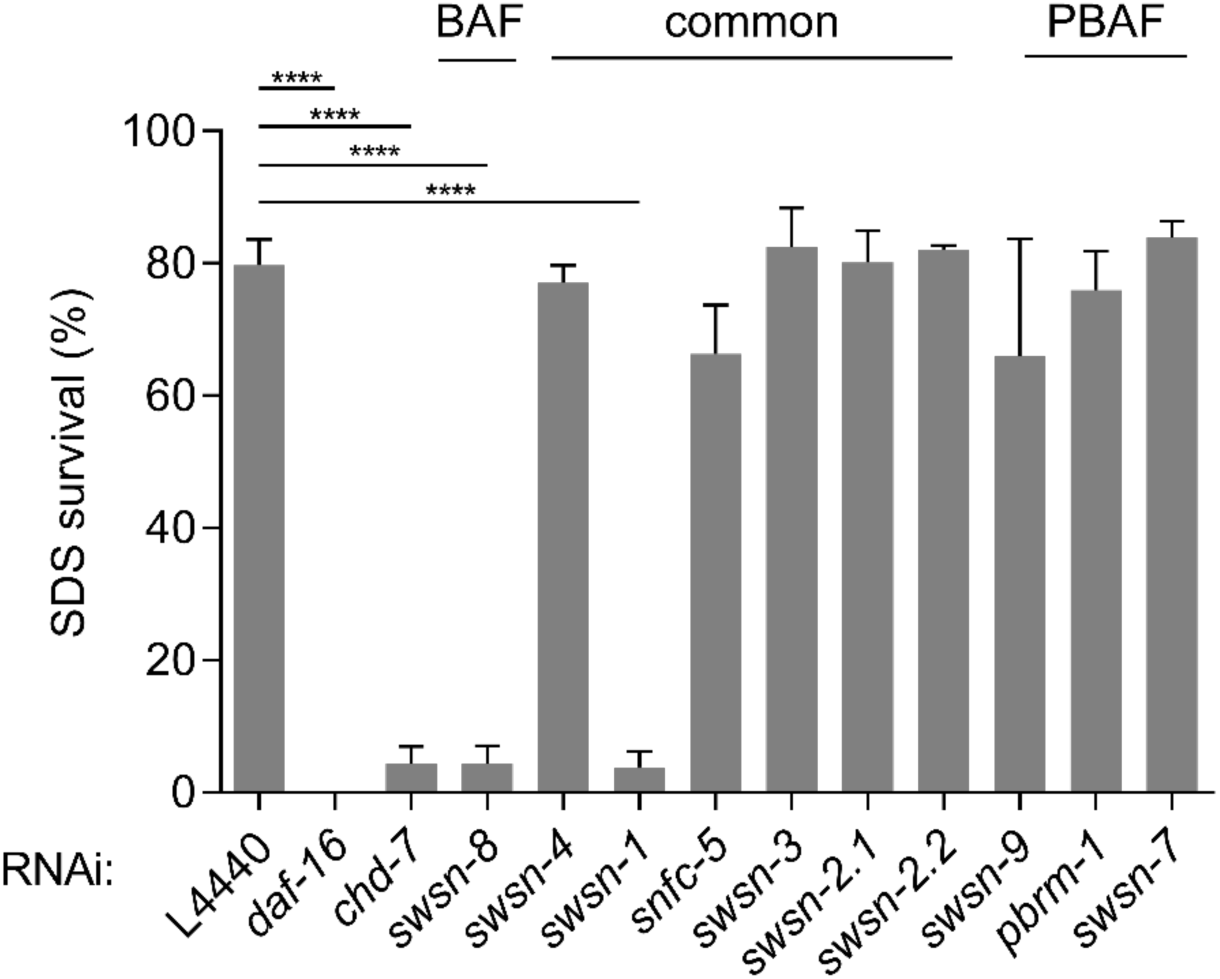
*swsn-*1 *and swsn-*8 share dauer suppression phenotypes with *chd-7*. *daf-7(e1372)* L1 synchronized larvae were placed on the indicated RNAi bacteria and grown at 25°C for 4 days to induce dauer formation. Dauer were identified after 4-5 days based on morphology and their resistance to 1% SDS for 15 min. Common SWI/SNF subunits and BAF or PBAF subclasses are indicated on top. Lines above bars represent standard error of the mean (SEM) from three independent experiments. At least 60 animals were assayed for each RNAi. One-way ANOVA. ****p <0.0001. *comparison to the control, L4440 RNAi.

**Figure S4:**
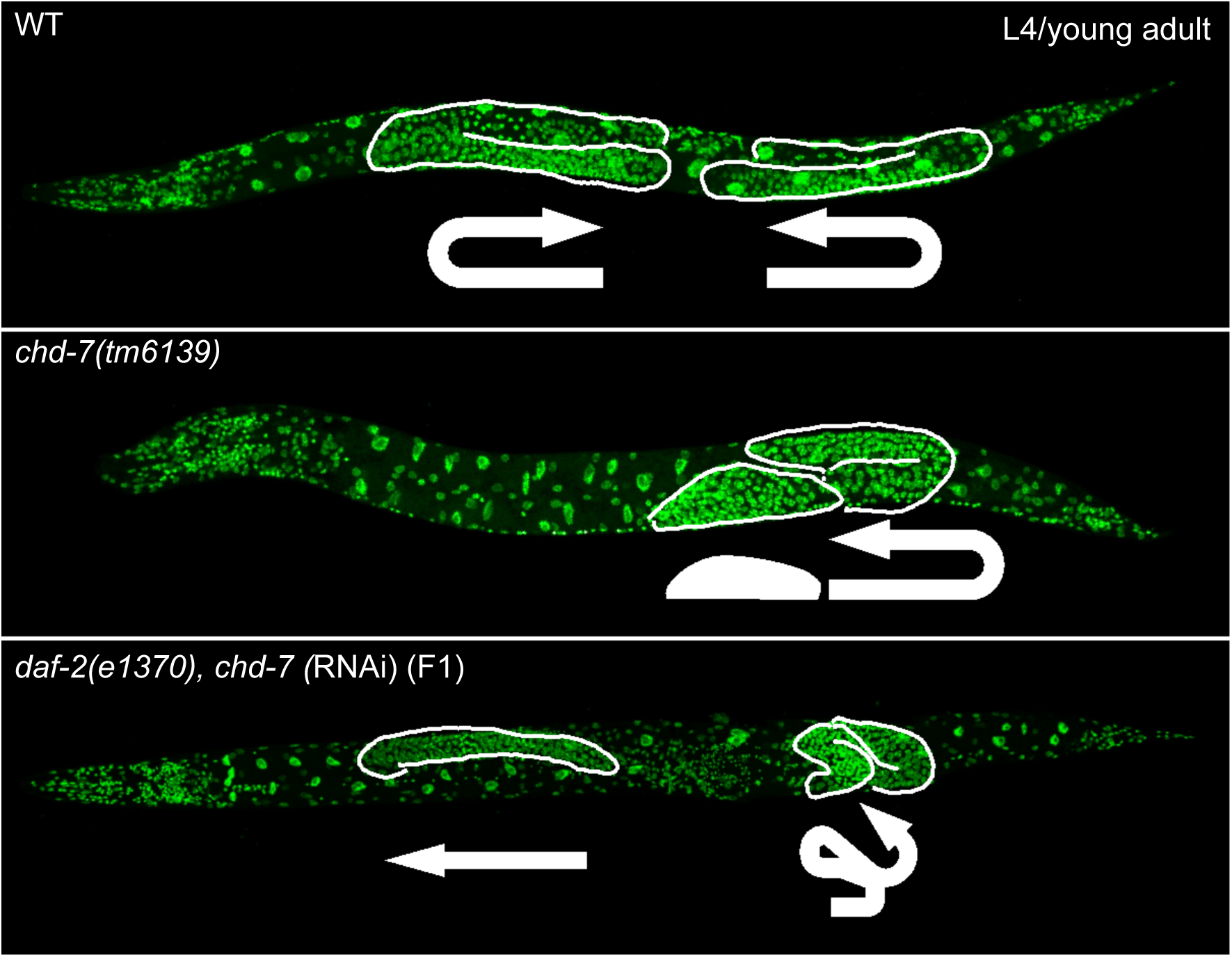
c*hd-7* mutants show altered gonad proliferation and migration. Whole mount fixation and DAPI (green) staining of late L4/young adult animals from the relevant strains. RNAi sample was obtained by growing *daf-2(e1370)* for two generations on *chd-7* dsRNA-producing E.Coli..

**Figure S5:**
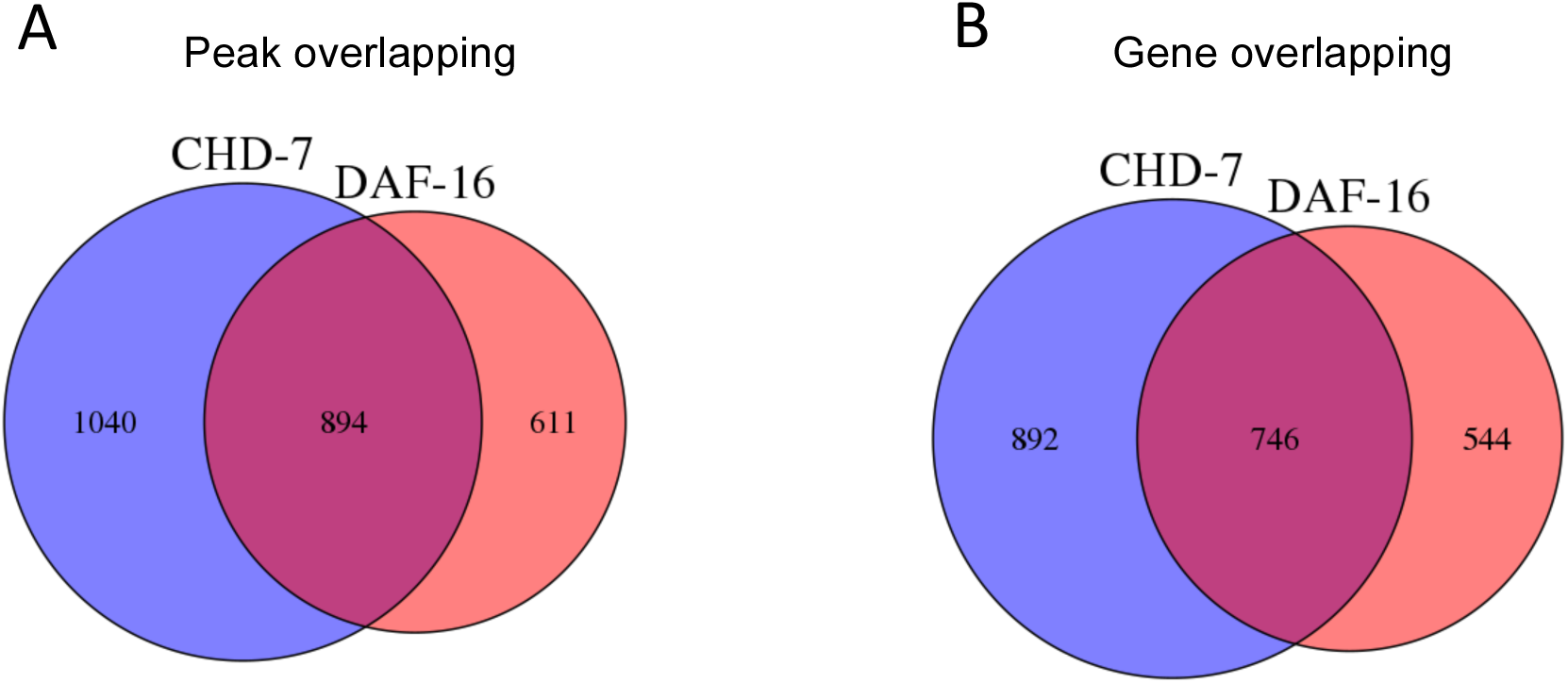
ChIP-seq analysis of CHD-7 and DAF-16 from ModEncode. Comparison of gene lists from peak calling of CHD-7::GFP (young adult) and DAF-16::GFP (L4) using Homer software. A) Venn diagram of ChIP peaks for CHD-7 and DAF-16. Overlap was defined as sharing at least one base in common. B) Common genes shared by CHD-7::GFP and DAF-16::GFP peaks. CHIP-seq data was obtained from publicly available data from the ModEncode project.

**Figure S6:**
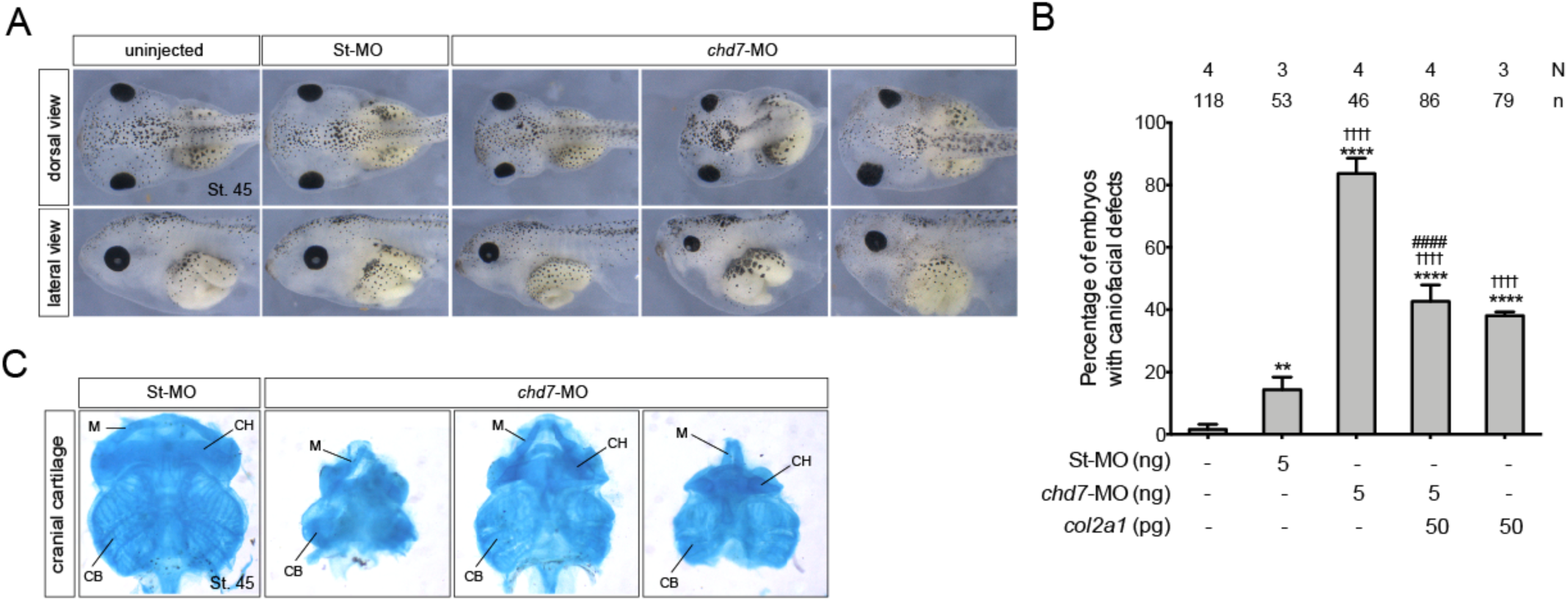
Craniofacial defects of *Chd7* knock-down *Xenopus* tadpoles. A) Gross morphology of surviving tadpoles at St. 45. Embryos were injected into both D1 blastomeres at the 8-cell stage with St-MO (5 ng) and *chd7*-MO (5 ng) as indicated. Tadpoles position is anterior to the left. Embryos are representative. B) Quantification of *Xenopus* tadpoles with craniofacial defects at stage 45. Embryos were injected as indicated in A and in the graph. Data on graph is presented as means with standard error. Fisher’s exact test (\*\**p*< 0.01, ****^,††††,####^*p*< 0.0001). *comparison to uninjected group, ^†^ comparison to St-MO group, ^#^ comparison to *chd7*-MO group. N = number of experiments, n = number of embryos. C) Examples of Alcian blue-stained craniofacial skeletal elements from St. 45 tadpoles injected with St-MO (5 ng) and *chd7*-MO (5 ng). M: Meckel’s, CH: ceratohyle and CB: ceratobranchial cartilages. *chd7*-MO injected tadpoles presented a high incidence of gross head malformations, reduced head and eyes sizes, and abnormal development of craniofacial cartilages.

